# Computational insights into differential interaction of mamalian ACE2 with the SARS-CoV-2 spike receptor binding domain

**DOI:** 10.1101/2021.02.02.429327

**Authors:** Cecylia S. Lupala, Vikash Kumar, Xiao-dong Su, Chun Wu, Haiguang Liu

## Abstract

The severe acute respiratory syndrome coronavirus 2 (SARS-CoV-2), the causing agent of the COVID-19 pandemic, has spread globally. Angiotensin-converting enzyme 2 (ACE2) has been identified as the host cell receptor that binds to receptor-binding domain (RBD) of the SARS-COV-2 spike protein and mediates cell entry. Because the ACE2 proteins are widely available in mammals, it is important to investigate the interactions between the RBD and the ACE2 of other mammals. Here we analyzed the sequences of ACE2 proteins from 16 mammals and predicted the structures of ACE2-RBD complexes. Analyses on sequence, structure, and dynamics synergistically provide valuable insights into the interactions between ACE2 and RBD. The comparison results suggest that the ACE2 of bovine, cat and panda form strong binding with RBD, while in the cases of rat, least horseshoe bat, horse, pig, mouse and civet, the ACE2 proteins interact weakly with RBD.

## Introduction

The severe acute respiratory syndrome coronavirus 2 (SARS-CoV-2), a novel coronavirus, is responsible for the new type of severe pneumonia COVID-19 [1]. As of December 6, 2020, over 66 million people have been tested positive for the SARS-CoV-2, and the number of infections still rapidly increases [2]. SARS-CoV-2 utilizes the human angiotensin-converting enzyme 2 protein (ACE2) to initiate the spike protein binding and to facilitate the viral attachment to host cells[3–8]. Recently, reports of other animals testing positive for SARS-CoV-2 are emerging. Studies on viral replication and susceptibility to SARS-CoV-2 suggested that the virus replicates efficiently in cats or ferrets [9]. There are reports of dog, cat and tiger testing positive for SARS-CoV-2[10–12]. Therefore, it is highly desirable to study the susceptibility of those mammalian animals, which are in close contact with humans. Because ACE2 proteins exist in many mammailian animals, potentially making them susceptible to SARS-CoV-2, we gathered ACE2 sequences of 16 animals for detailed analysis (See **Table 1**). By studying the interactions between the receptor binding domain (RBD) of virus spike protein and ACE2 receptors, we hope to provide information on animal susceptibility to the SARS-CoV-2. It has been established that the RBD of the SARS-CoV-2 (denoted as RBD hereafter) and the human ACE2 (hACE2) form stable complexes, as shown in recently determined crystal structures[13,14] and computer simulations[15]. This provides an opportunity to investigate the interactions between RBD and ACE2 of other mammalian animals. Although such knowledge alone is not sufficient to accurately predict the susceptibility of animals to SARS-CoV-2, the information is valuable in understanding the interactions between RBD and ACE2.

**Table 1.**
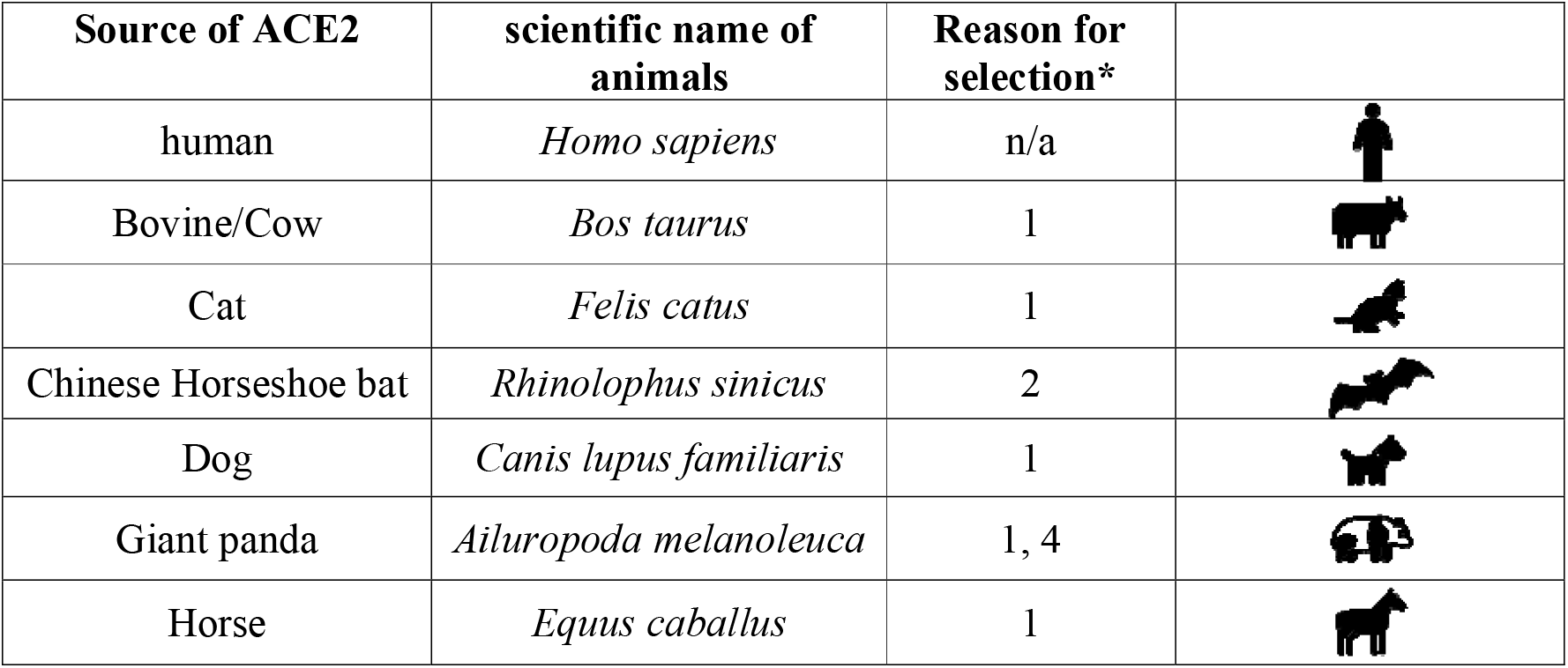
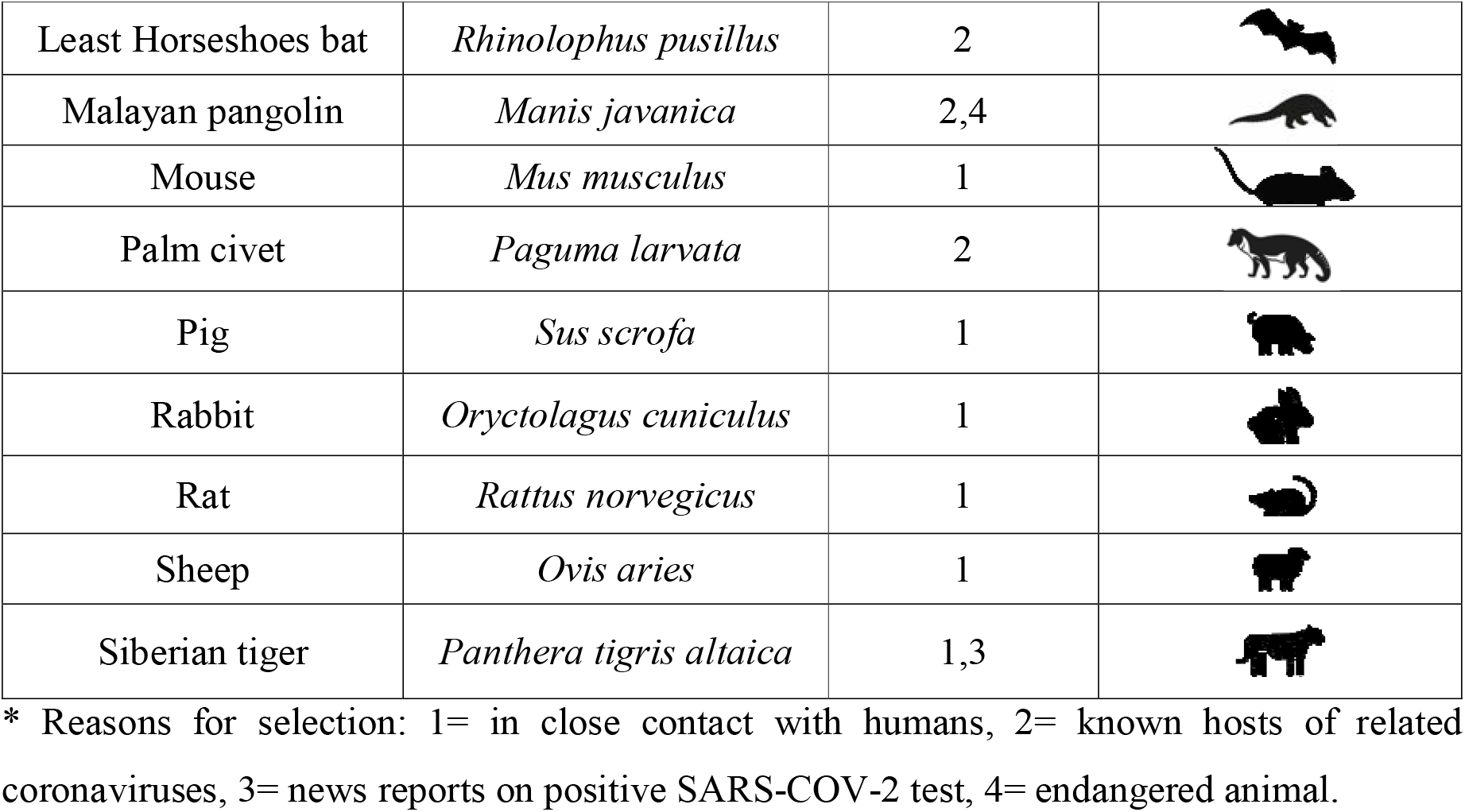
ACE2 proteins selected in this study.

| Source of ACE2 | scientific name of animals | Reason for selection* |
| --- | --- | --- |
| human | *Homo sapiens* | n/a |
| Bovine/Cow | *Bos taurus* | 1 |
| Cat | *Felis catus* | 1 |
| Chinese Horseshoe bat | *Rhinolophus sinicus* | 2 |
| Dog | *Canis lupus familiaris* | 1 |
| Giant panda | *Ailuropoda melanoleuca* | 1, 4 |
| Horse | *Equus caballus* | 1 |
| Least Horseshoes bat | *Rhinolophus pusillus* | 2 |
| Malayan pangolin | *Manis javanica* | 2,4 |
| Mouse | *Mus musculus* | 1 |
| Palm civet | *Paguma larvata* | 2 |
| Pig | *Sus scrofa* | 1 |
| Rabbit | *Oryctolagus cuniculus* | 1 |
| Rat | *Rattus norvegicus* | 1 |
| Sheep | *Ovis aries* | 1 |
| Siberian tiger | *Panthera tigris altaica* | 1,3 |

\* Reasons for selection: 1= in close contact with humans, 2= known hosts of related coronaviruses, 3= news reports on positive SARS-COV-2 test, 4= endangered animal.

The conservation of ACE2 residues and structures of ACE2-RBD complexes are reported in a few studies [16–18], dynamics simulations were also applied to investigate the dynamical features of the ACE2-RBD interactions[19,20]. In this report, we combined sequence analysis, structure prediction, molecular dynamics to investigate the interactions between ACE2 and RBD. Using the crystal structure of SARS-CoV-2 RBD and human ACE2 complex (hACE2-RBD) as the template[21], ACE2-RBD complex structures were constructed for previously mentioned ACE2 proteins, and the dynamics of these complexes were investigated using simulations. Based on conservation in ACE2 residues, similarity in electrostatic potentials, and dynamical interactions revealed from simulations, we classified these ACE2-RBD interactions into weak, medium, and strong categories.

## Materials and Methods

### Homology modeling of the ACE2-RBD complex structures

The ACE2 sequences were obtained from the NCBI and uniport databases[22,23]. Using the SWISS-MODEL interactive server[24], we modeled structures for 15 mammalian ACE2 proteins, based on hACE2 structure (PDB ID: 6LZG[21]). Then, the SARS-CoV-2 RBD and ACE2 complexes were assembled by superposing the homology structure of ACE2 proteins to hACE2-RBD complex structure.

### Comparison of the electrostatic potential of ACE2 on RBD binding interface

PyMOL (https://pymol.org/2/) was utilized to compute and visualize the electrostatic potential maps at the ACE2-RBD complex interfaces. These maps depict the electrostatic potential surface rendered from the numerical solutions of the Poisson-Boltzmann equation[25]. The electrostatic potential surfaces were simplified into 2D projection images for pairwise comparison and clustering analysis. The hierarchical clustering algorithm was applied to group these 2D projection images.

### Molecular dynamics simulations of ACE2-RBD complexes

GROMACS-5.1.2[26] was used to carry out MD simulations of ACE2-RBD complexes. All complexes were parametrized with CHARMM27 force fields [27]. Disulfide bonds were maintained as in the crystal structure of hACE2-RBD complex. Complexes were solvated in the triclinic box with a minimum distance of 10 Å between the complex and the box boundaries. Solvated systems were neutralized by adding ions (Na^+^ and Cl^-^) to 0.15 mM. Then, these systems were subjected to steepest descent energy minimization, followed by constant volume (NVT) and constant pressure of 1 bar (NPT) equilibrations, for 1 ns and 3 ns respectively. During system equilibration, positional restraints were applied on non-hydrogen atoms of ACE2-RBD complexes. Temperature and pressure were controlled by the V-rescale method[28] and Parrinello-Rahman method[29], respectively. Finally, 50 ns production simulations were carried out at NPT condition. VMD[30] and UCSF Chimera[31] were used to visualize and analyze simulation trajectories. The physical binding interactions comprising van der Waals and electrostatic components were calculated between each ACE2 and RBD for the structures sampled in MD simulations.

## Results and Discussions

### Sequence analysis and the conservation at the RBD binding interface

Multiple sequence alignment was carried out using CLUSTALW program[32], [33], and the aligned sequences were redrawn with the human ACE2 crystal structure as the reference using the ESPript webserver[34]. All ACE2 proteins comprise amino acids from position 19 to 614, except for dog ACE2, which has a deletion at position 20 (**Figure 1 and Figure S1)**. In the human ACE2-RBD complex, the amino acids of ACE2 at the N-terminal helix-1 (residues 19-42), near the η1 (residues 82-83), helix-13 (residue 330) and β-hairpin-4,5 (residues 352-357), have been identified as the key residues (**Figure 1**) that bind to the RBD [13,15].

**Figure 1.**
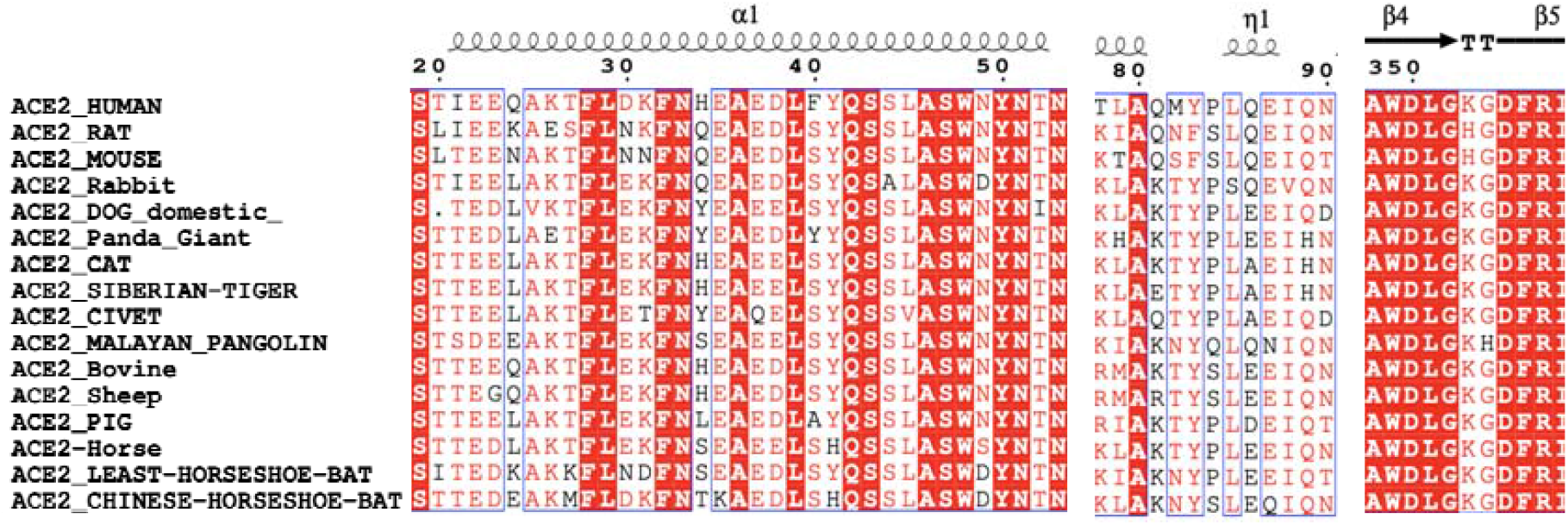
The comparison for the key residues at the binding interfaces after multiple sequence alignment analysis.

Based on the hACE2-RBD structure, we further identified 13 key residues on the RBD-interacting interface and analyzed their sequence conservations comparing to hACE2 (**Figure 2**). Siberian tiger and cat share the same ACE2 protein residues among these 13 residues, so the analysis on tiger ACE2 is inferred from cat ACE2. Taking hACE2 sequence as the reference, substituted residues of these 13 positions are summarized in **Figure 2A and Table S1**. We found that residues at positions 24, 27, 31, 34 and 82 are highly variable among these ACE2 proteins. The H34 of hACE2 has the largest variation, which is substituted by Q (Rat, Mouse and Rabbit), Y (Dog, Giant Panda and Civet), S (Pangolin, Horse and Least Horseshoe Bat), L (Pig), and T (Chinese Horseshoe Bat). The Q24 has four variations: K (Rat and Least Horseshoe Bat), N (Mouse), L (Rabbit, Dog, Cat, Giant Panda, Civet, Pangolin, Pig and Horse), and E (Chinese Horseshoe Bat). T27, K31, and M82 all have three different substitutions, while D30 and M82 have two different possible substitutions. **Figure 2B** shows the number of identical residues to hACE2 at these 13 positions. Bovine and sheep ACE2 proteins differ from hACE2 at only two positions, while the ACE2 proteins of mouse, rat and civet are different from hACE2 at 7 out of 13 positions.

**Figure 2.**
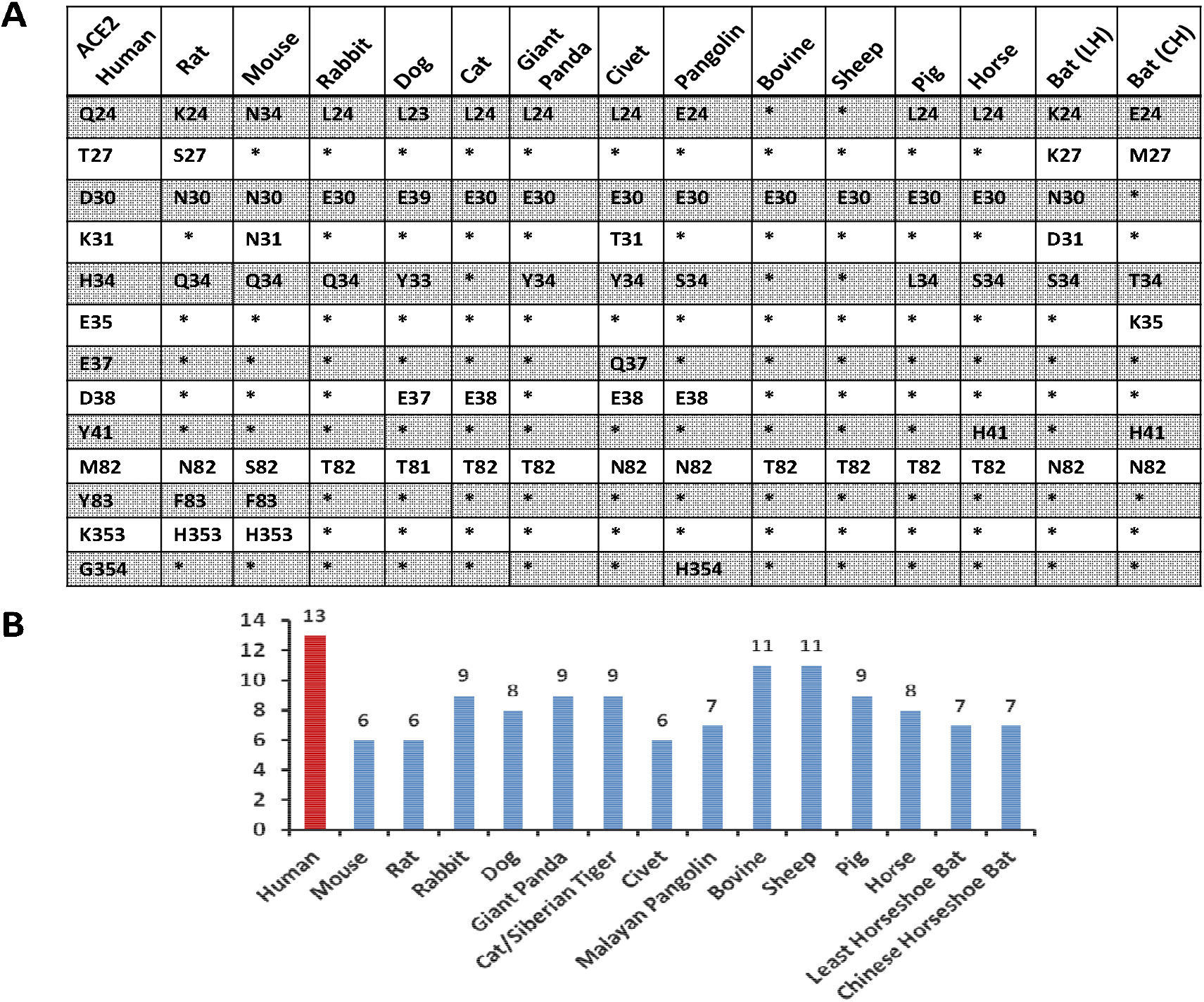
Residue conservation analysis. **(A)** Comparison of 13 critical residues in binding to SARS-CoV-2 RBD. Bat (LH) stands for Least Horseshoe and Bat (CH) stands for Chinese Horseshoe Bat. Cat ACE2 is used to represent both cat and Siberian tiger ACE2 proteins, their sequences are identical at these 13 positions. (**B**) Number of identical residues compared to hACE2 at the 13 positions marked in (A).

### Electrostatic potential surface at the binding interface

The electrostatic potential surfaces for the central region of ACE2 helix-1 (residues 30-37) are shown in **Figure 3** for all ACE2-RBD complexes. According to electrostatic potential maps, this region features a charge distribution composed of both positively and negatively charged sites in human, bovine and cat ACE2, while the electrostatic potentials are mostly negative for dog and pig ACE2 proteins. Clustering analysis on electrostatic potential surfaces showed that the bovine/sheep/pig/rabbit ACE2 proteins have similar features as hACE2 in this region (**Figure 3C**). The mouse/rate/least-horseshoe-bat ACE2 show the least similarity in the electrostatic potential features in this region comparing to other ACE2 proteins.

**Figure 3.**
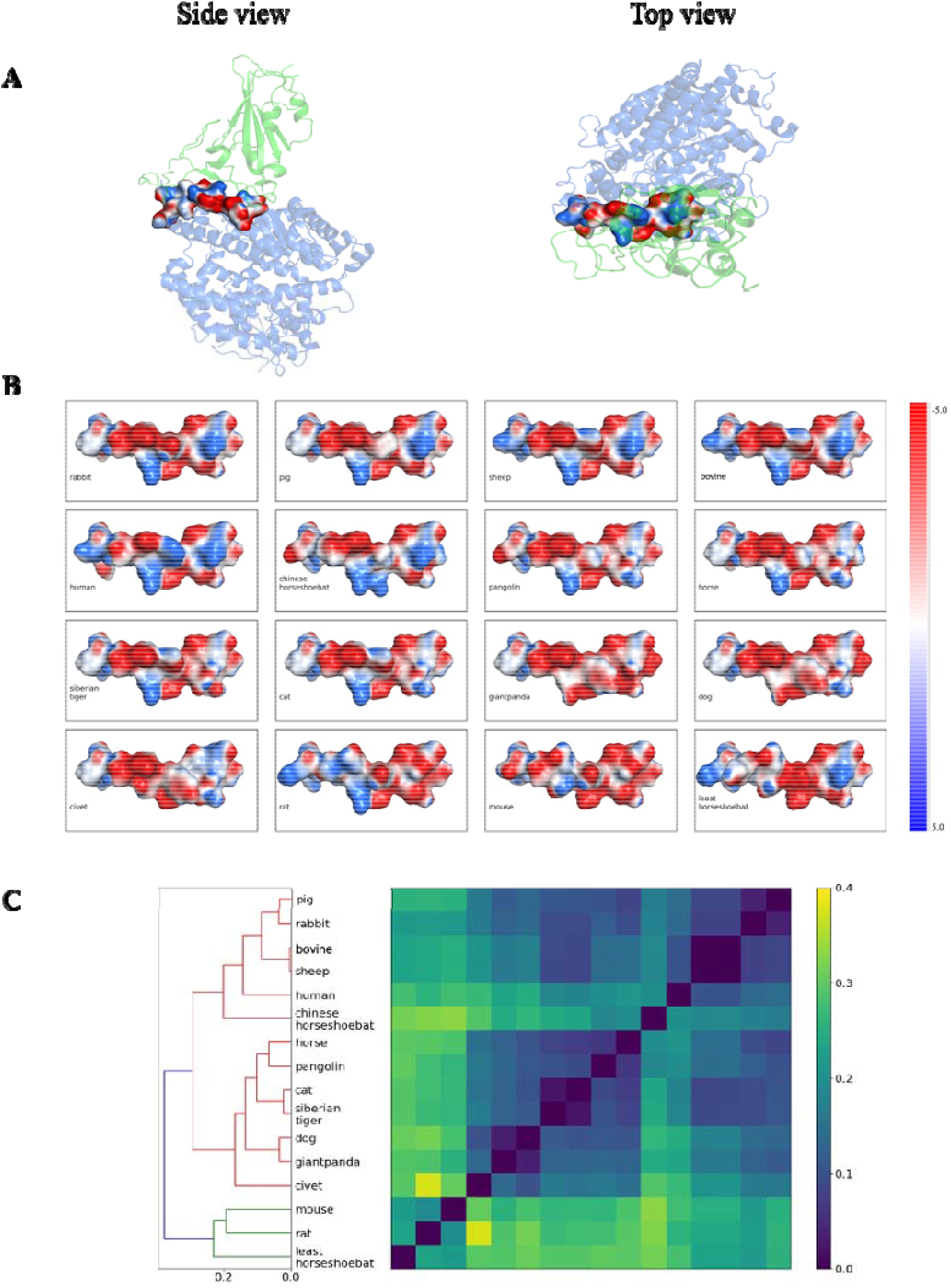
Electrostatic potential surface analysis. **(A)** ACE2 binding interface to RBD at two orientations. **(B)** The top view of the electrostatic potential surfaces for central binding region of between ACE2 and SARS-CoV-2-RBD. In human, cat and bovine ACE2, positions 30-37 comprise both positive and negatively charged residues. The residue substitutions in ACE2 of dog and civet at the same region lead to negatively charged patches. (**C**) Hierarchical clustering results based on similarity between electrostatic potential surfaces

### Binding interactions assessed from MD simulation data

Simulation trajectories of 16 ACE2-RBD complexes were analyzed with a focus on the ACE2 residues at the binding interface (**Figure S2-S4**). Considering the critical roles of hydrogen bonding interactions between ACE2 and RBD, we focused on the analysis of interfacing hydrogen bonds. The occupancies of hydrogen bonds were calculated from simulation trajectories (**Figure 4** and **Figure S5)**. For these ACE2-RBD complexes, five frequently observed hydrogen bonds are D30:K417, E35:Q493, Y83:N487, K353:G502 and D355:T500 (ACE2 residues are placed on the left of colon, and RBD residues on the right). In the following, detailed discussions are grouped based on the number of substitutions among the 13 key residues.

**Figure 4.**
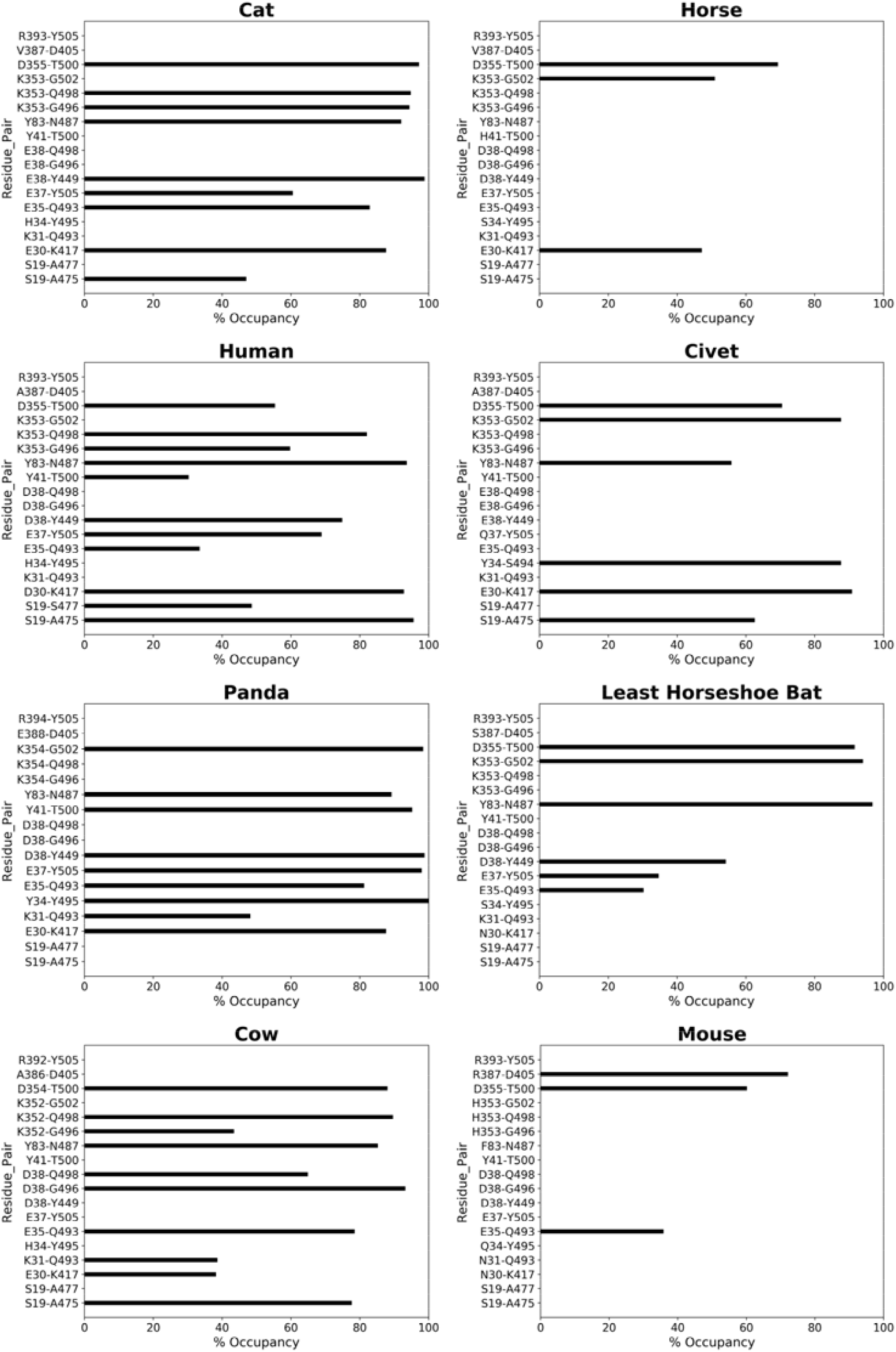
Occupancies of hydrogen bonds at ACE2-RBD interface. The left panels are hydrogen bonding patterns in strong binding cases (as labeled above each plot); the right panels correspond to the weak binding cases. Each hydrogen bond comprises one residue from ACE2 and one from RBD, shown on the left and right of the hyphen respectively.

Bovine ACE2-RBD shows a highly similar hydrogen bonding pattern as hACE2-RBD (**Figure 4**). Cat ACE2 shows even stronger hydrogen bonding interactions with RBD than hACE2, but the hydrogen bond (H-bond) between Y41 and T500 is absent (**Figure 4**). Cat ACE2-RBD also exhibits the highest interaction energy among 16 ACE2-RBD complexes (see **Table 2**). This is consistent with recent reports on domestic cat being infected by SARS-CoV-2[9,10]. A recently resoved complex structure of cat ACE2-RBD reveals similar binding as the hACE2-RBD complex[35]. Experimental data also show that cats are efficient in replicating SARS-CoV-2 [9], suggesting that four substitutions do not inhibit RBD binding.

**Table 2.** Molecular interaction energies between ACE2 and RBD. 250 structures from the simulations were used to compute interaction energies. Strong interactions are highlighted in bold font.

| Complex | Interaction energy of homology models (kcal/mol) | No. of H-bonds having >30 % occupancy | Average interaction energy (kcal/mol) |
| --- | --- | --- | --- |
| **Human** | **-169.56** | **11** | **-170.08±10.79** |
| Rat | -135.33 | 5 | -125.69±15.52 |
| Mouse | -127.39 | 3 | -107.31±9.16 |
| Rabbit | -165.99 | 9 | -151.48±15.56 |
| Dog | -168.71 | 7 | -147.79±14.23 |
| Siberian Tiger | -155.13 | 7 | -144.82±13.43 |
| **Cat** | **-170.80** | **9** | **-175.77±12.20** |
| Civet | -155.95 | 6 | -115.67±12.16 |
| **Bovine** | **-182.96** | **10** | **-162.83±14.04** |
| Least Horseshoe Bat | -157.98 | 6 | -114.12±17.08 |
| Malayan Pangolin | -179.45 | 9 | -155.46±10.94 |
| Pig | -172.54 | 4 | -127.67±13.74 |
| Chinese Horseshoe Bat | -176.70 | 5 | -140.40±13.55 |
| Horse | -148.84 | 3 | -119.62±11.46 |
| Sheep | -181.49 | 9 | -146.63±15.32 |
| **Panda** | **-187.17** | **9** | **-167.30±9.92** |

Panda and pig ACE2 proteins both differ from hACE2 at four positions, but their interactions with RBD are quite different. Panda ACE2 forms 9 strong hydrogen bonds with RBD (**Figure 4**). The pig ACE2 interacts with RBD more weakly than panda ACE2, in line with experimental studies showing that SARS-CoV-2 infection was not detectable in pigs or their cell lines[36]. The difference in the interaction profiles of panda and pig may be due to Y34L substitution and the influences from other domains of ACE2.

Dog and horse ACE2 have five substitutions (four are at positions 24,30,34,82; and one occurs at position 38 for dog and position 41 for horse). Dog is the first domestic animal reported testing positive with a low level of SARS-CoV-2 infection[37,38]. The dog ACE2 contains a deletion at position 20 (threonine in human ACE2), but this deletion does not appear to affect the complex structures revealed in the homology models. In the case of dog ACE2-RBD (**Figure S3**), S19, E37, Y40, Q41 do not form hydrogen bonds with RBD (**Figure S5**), weakening the interactions of dog ACE2 with RBD. Horse ACE2 forms only 3 hydrogen bonds with RBD (**Figure 4**).

With respect to hACE2, pangolin, CH-bat and LH-bat ACE2 differ at 6 positions. Due to the co-evolution with other coronaviruses, pangolin and bats were speculated to be intermediate hosts of SARS-CoV-2. Despite having 6 substitutions, pangolin ACE2 forms strong interactions with RBD. Previously, Q24K mutant of ACE2 revealed a slight inhibition effect on the binding to the RBD of SARS-CoV spike protein, and the binding is abolished for K31D mutant[39]. The high similarity between RBD of the two coronaviruses suggest that the mutant may exert a similar effect to SARS-CoV-2 infection. LH-bat ACE2 has both substitutions, leading to weak interactions with only 6 hydrogen bonds (**Figure 4**).

Civet, mouse, and rat have 7 substitutions in their ACE2 compared to hACE2. Civets can be infected by coronaviruses in natural environments [40]. In civet ACE2, the important hydrogen bond D30:K417 in hACE2-RBD is not formed between E30 of civet ACE2 and K417 of RBD (**Figure 4**). Mouse ACE2 shows the weakest interaction with RBD, with only 3 hydrogen bonds. The mutation Y83F in both mouse/rat results in the loss of the hydroxyl moiety of tyrosine (in hACE2), losing a hydrogen bond with the N487 of the RBD. This Y83F mutation has been reported to inhibit interaction with SARS-CoV spike RBD[39]. Another noticeable substitution occurs at the highly conserved K353, which is replaced by histidine in both mouse and rat ACE2 (**Figure 2**). The K353H substitution eliminated hydrogen bonds with N501 of RBD, exerting a significiant impact on RBD binding.

The interaction between RBD and ACE2 is quantified by molecular mechanics energy comprising van der Waals and electrostatic interaction terms. As summarized in **Table 2**, the ACE2 of cat, panda, bovine, and human form strong interactions with the RBD, while the ACE2-RBD interactions are much weaker in the cases of mouse, least horseshoe bat, civet, horse, rat, and pig. The sequence conservation, molecular interactions are correlated to experimental results (Table 3), providing insights on the interactions at molecular levels.

**Table 3.** Binding interactions classified based on sequence identity and interactions.

| Mammals | Experimental results | Sequence identity | Interaction energy |
| --- | --- | --- | --- |
| Human | High [1,41–43] | High | High |
| Bovine | Not available | High (11/13) | High |
| Sheep | Not available | High (11/13) | Medium |
| Cat | High [9,10,44] | Medium (9/13) | High |
| Tiger | Medium [12] | Medium (9/13) | Medium |
| Panda | Not available | Medium (9/13) | High |
| Pig | Not susceptible [36] | Medium (9/13) | Low |
| Rabbit | Low [44] | Medium (9/13) | Medium |
| Dog | Medium [9,44] | Medium (8/13) | Medium |
| Horse | Not available | Medium (8/13) | Low |
| Pangolin | Medium [44] | Medium (7/13) | Medium |
| Bat (LH) | Not available | Medium (7/13) | Low |
| Bat (CH) | Medium [45] | Medium (7/13) | Medium |
| Civet | Not susceptible [44] | Low (6/13) | Low |
| Mouse | Not susceptible [8,44] | Low (6/13) | Low |
| Rat | Not susceptible [44] | Low (6/13) | Low |

Classification criteria: Sequence identity of 13 residues compared to hACE2 (Low, 0-50%; Medium, 50-75%; High, 75%-100%); Interaction energy (Low, -100 ~ -130 kcal/mol; Medium, -130 ~ -160 kcal/mol; and High, -160 ~ -190 kcal/mol).

## Conclusions

Sequence, structure, and dynamics are analyzed for 16 ACE2 proteins in complex with the RBD of SARS-CoV-2 spike protein. The ACE2 of bovine and sheep exhibit high sequence identities to human ACE2. MD simulations reveals that bovine, cat and panda ACE2 proteins show strong binding interactions with the RBD. ACE2 of dog, siberian tiger, malayan pangolin, rabbit, sheep and rabbit show relatively weaker interactions. This study provides a molecular basis for differential interactions between ACE2 and RBD in 16 mammals and will be useful in predicting the host range of the SARS-CoV-2.

## Acknowledgement

The work is supported by Beijing Computational Science Research Center (CSRC) via a director discretionary grant. The research is supported by National natural science foundation (NSFC Grant number: U1930402).

