## supporting information for "Computational insights into differential interaction of mamalian ACE2 with the SARS-CoV-2 spike receptor binding domain"

**Table S1: Details of amino acid substitutions compared to human ACE2.**

| **Source of ACE2**  **(**number of substitutions**)** | **Substitution (change in amino acid type)** |
| --- | --- |
| Rat (7) | Q24K (**Polar to positively charged**)  T27S (Polar to polar)  D30N (**Negatively charged to polar**)  H34Q (Polar to polar)  M82N (**Non-polar to polar)**  Y83F (**Polar to non-polar**)  K353H (**Positively charged to polar**) |
| Mouse (7) | Q24N (Polar to polar)  D30N (**Positively charged to polar**)  K31N (**Negatively charged to polar**)  H34Q (Polar to polar)  M82S (**Non-polar to polar**)  Y83F (**Polar to non-polar**)  K353H (**Positively charged to polar**) |
| Rabbit(4) | Q24L(**Polar to non-polar**)  D30E(Negatively charged to negatively charged)  H34Q(Polar to polar)  M82T (**Non-polar to polar**) |
| Dog(5) | Q24L(**Polar to non-polar**)  D30E(Negatively charged to negatively charged)  H34Y(Polar to polar)  D38E(Negatively charged to negatively charged)  M82T(**Non-polar to polar**) |
| Giant Panda(4) | Q24L(**Polar to non-polar**)  D30E(Negatively charged to negatively charged)  H34Y(Polar to polar)  M82T(**Non-polar to polar**) |
| Cat(4) | Q24L(**Polar to non-polar**)  D30E(Negatively charged to negatively charged)  D38E (Negatively charged to negatively charged)  M82T(**Non-polar to polar)** |
| Siberian Tiger (4) | Q24L(**Polar to non-polar**)  D30E(Negatively charged to negatively charged)  D38E (Negatively charged to negatively charged)  M82T(**Non-polar to polar**) |
| Civet(7) | Q24L(**Polar to non-polar**)  D30E(Negatively charged to negatively charged)  K31T(**Positively charged to polar**)  H34Y(Polar to polar)  E37Q(**Negatively charged to polar**)  D38E(Negatively charged to negatively charged)  M82T(**Non-polar to polar**) |
| Malayan Pangolin(6) | Q24E (**Polar to negatively charged**)  D30E (Negatively charged to negatively charged)  H34S (Polar to polar)  D38E (Negatively charged to negatively charged)  M82N(**Non-polar to polar**)  G354H (**Non-polar to polar**) |
| Bovine(2) | D30E(Negatively charged to negatively charged)  M82T(**Non-polar to polar**) |
| Sheep(2) | D30E(Negatively charged to negatively charged)  M82T(**Non-polar to polar**) |
| Pig(4) | Q24L(**Polar to non-polar**)  D30E(Negatively charged to negatively charged)  H34L(**Polar to non-polar**)  M82T(**Non-polar to polar**) |
| Horse(5) | Q24L(Polar to non-polar)  D30E(**Negatively charged to negatively charged**)  H34S(Polar to polar)  Y41H(Polar to polar)  M82T(**Non-polar to polar**) |
| Least Horseshoe Bat(6) | Q24K(**Polar to positively charged**)  T27K(**Polar to positively charged**)  D30N(**Negatively charged to polar**)  K31D(**Positively charged to negatively charged**)  H34S(Polar to polar)  M82N(**Non-polar to polar**) |
| Chinese Horseshoe Bat(6) | Q24E(**Polar to negatively charged**)  T27E(**Polar to negatively charged**)  H34T(Polar to polar)  E35K(**Negatively charged to positively charged**)  Y41H(Polar to polar)  M82T(**Non-polar to polar**) |

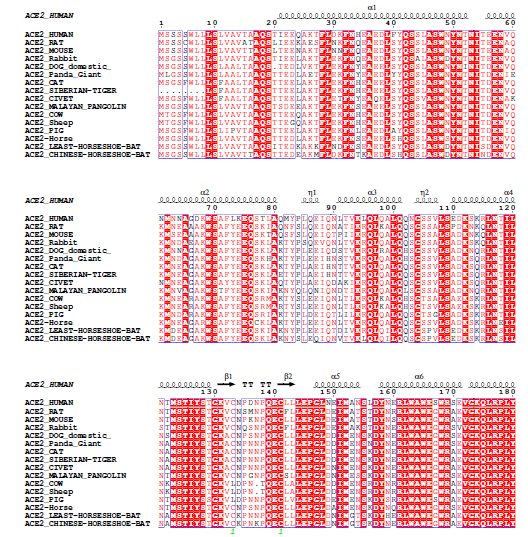

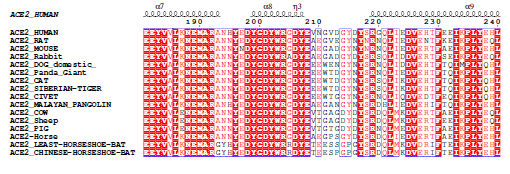

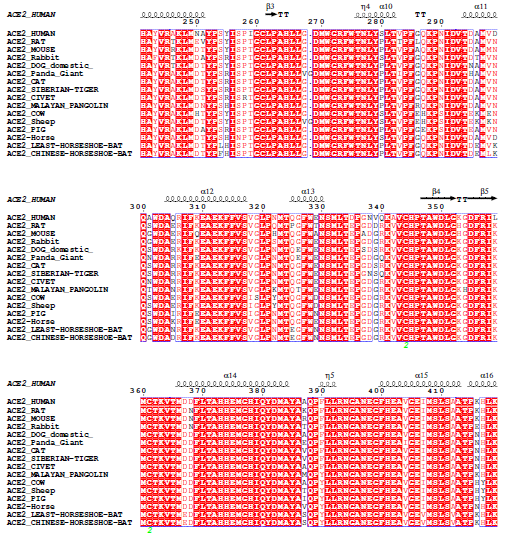

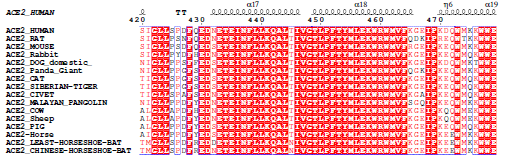

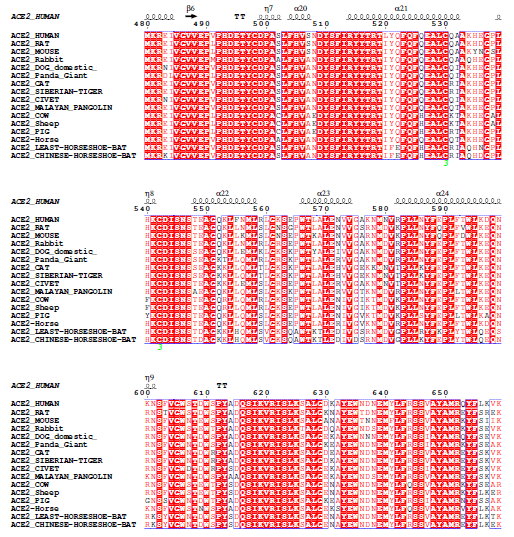

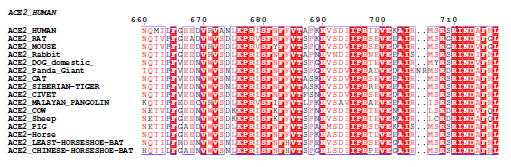

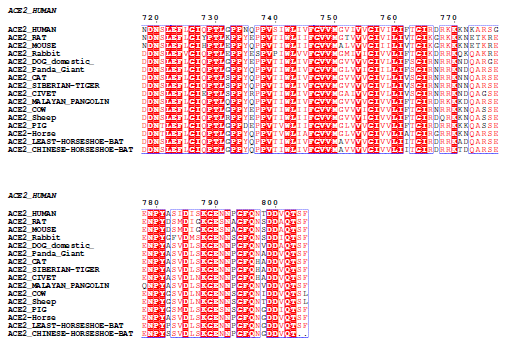

**Figure S1. Alignment of full ACE2 sequences of 16 mammals.**

**Sequences entries:**

Human (*Homo sapiens*) = [Q9BYF1](https://www.uniprot.org/uniprot/Q9BYF1) , rat (*Rattus norvegicus*) = [Q5EGZ1](https://www.uniprot.org/uniprot/Q5EGZ1) , mouse (*Mus musculus*) = [Q8R0I0](https://www.uniprot.org/uniprot/Q8R0I0) , rabbit (*Oryctolagus cuniculus*) = [G1TEF4](https://www.uniprot.org/uniprot/G1TEF4) , dog (*Canis lupus familiaris*) = [J9P7Y2](https://www.uniprot.org/uniprot/J9P7Y2) , giant panda (*Ailuropoda melanoleuca*) = [G1MC42](https://www.uniprot.org/uniprot/G1MC42) , cat (*Felis catus*) = [Q56H28](https://www.uniprot.org/uniprot/Q56H28) , siberian tiger (*Panthera tigris altaica*) = [XP_007090142.1](https://www.ncbi.nlm.nih.gov/protein/591328486) , civet (*Paguma larvata*) = [Q56N\]]-9L1](https://www.uniprot.org/uniprot/Q56NL1) , Malayan pangolin (*Manis javanica* ) = [XP_017505752.1](https://www.ncbi.nlm.nih.gov/protein/XP_017505752.1) ,Bovine (*Bos taurus*) = [Q58DD0](https://www.uniprot.org/uniprot/Q58DD0) , sheep (*Ovis aries*) = [W5PSB6](https://www.uniprot.org/uniprot/W5PSB6) , pig (*Sus scrofa*) = [K7GLM4](https://www.uniprot.org/uniprot/K7GLM4) , horse (*Equus caballus*) = XP_001490241.1 [F6V9L3](https://www.uniprot.org/uniprot/F6V9L3) , least horseshoe bat (*Rhinolophus pusillus*) = [E2DHI9](https://www.uniprot.org/uniprot/E2DHI9) and Chinese horseshoe bat (*Rhinolophus sinicus*) = [U5WHY8](https://www.uniprot.org/uniprot/U5WHY8)

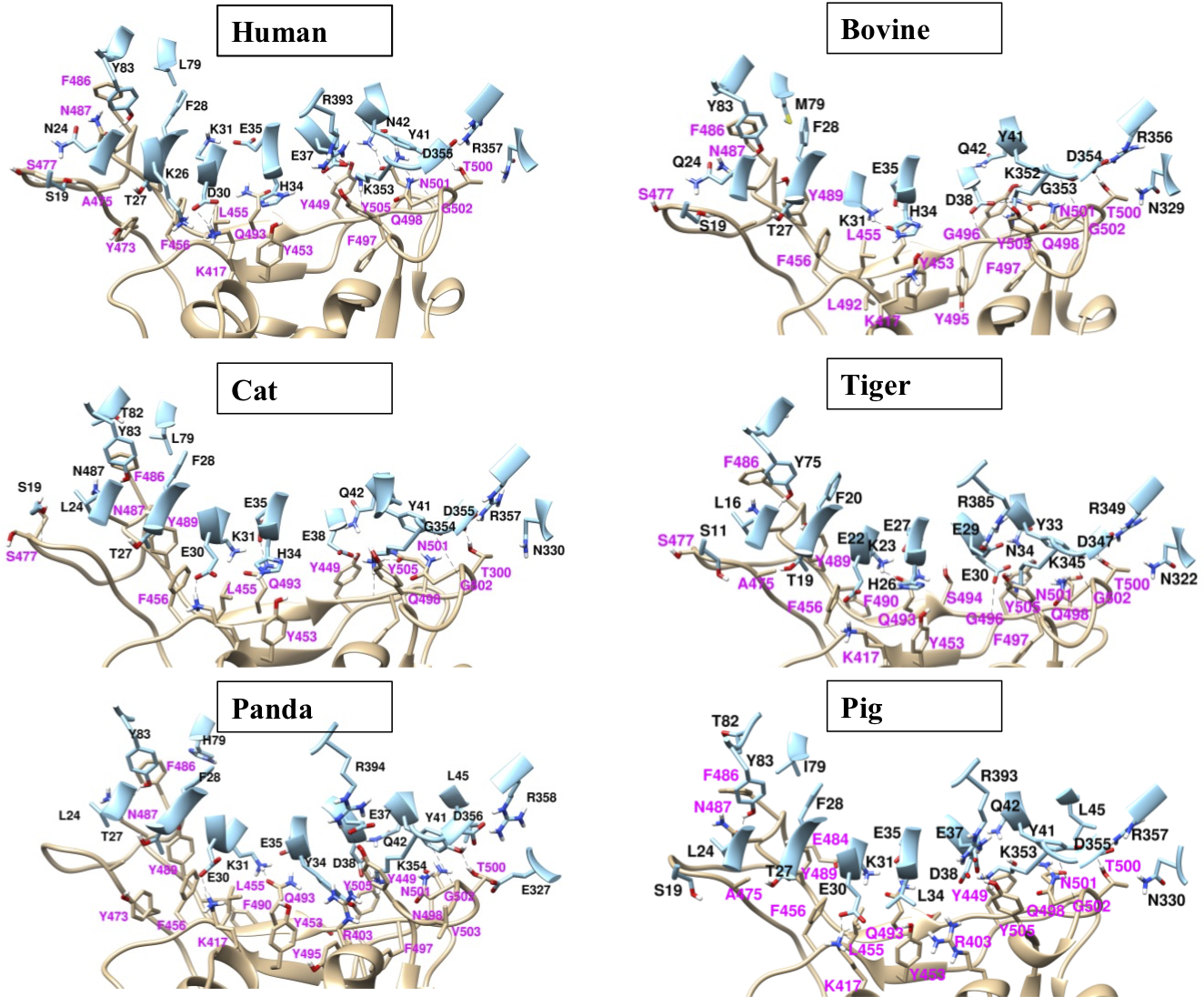

**Figure S2**. ACE2-RBD interface after 50 ns of MD simulations. ACE2 is shown in cyan color and SARS-CoV-2 spike in tan. Residues of ACE2 are labeled in black color, and the residues of RBD are labeled in pink color.

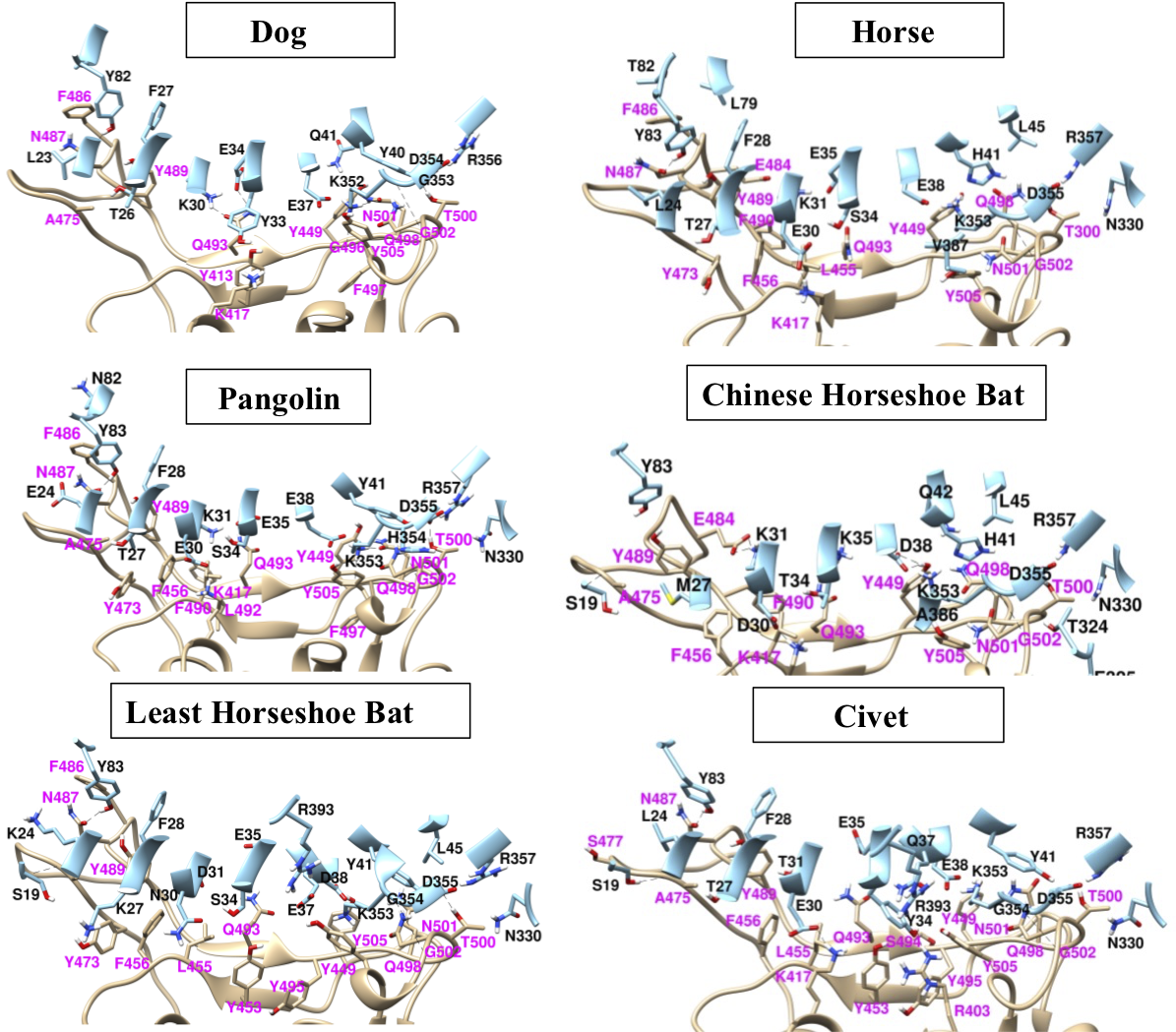

**Figure S3**. ACE2-RBD interface after 50 ns of MD simulations. ACE2 is shown in cyan color and SARS-CoV-2 spike in tan. Residues of ACE2 are labeled in black color, and the residues of RBD are labeled in pink color.

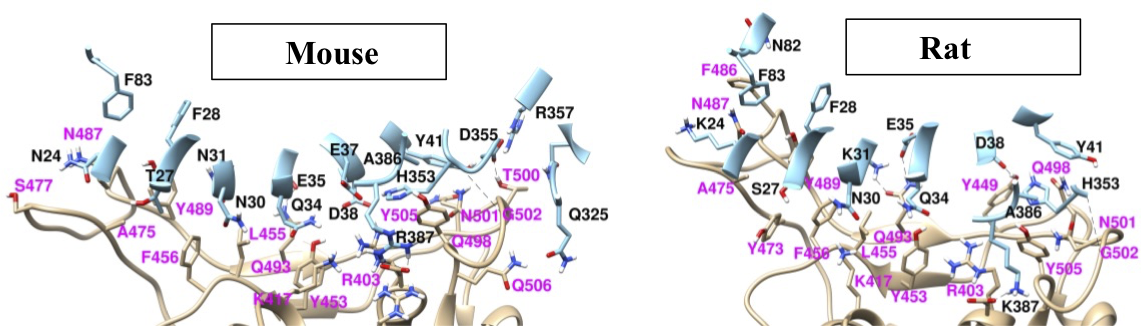

**Figure S4**. ACE2-RBD interface after 50 ns of MD simulations. ACE2 is shown in cyan color and SARS-CoV-2 spike in tan. Residues of ACE2 are labeled in black color, and the residues of RBD are labeled in pink color.

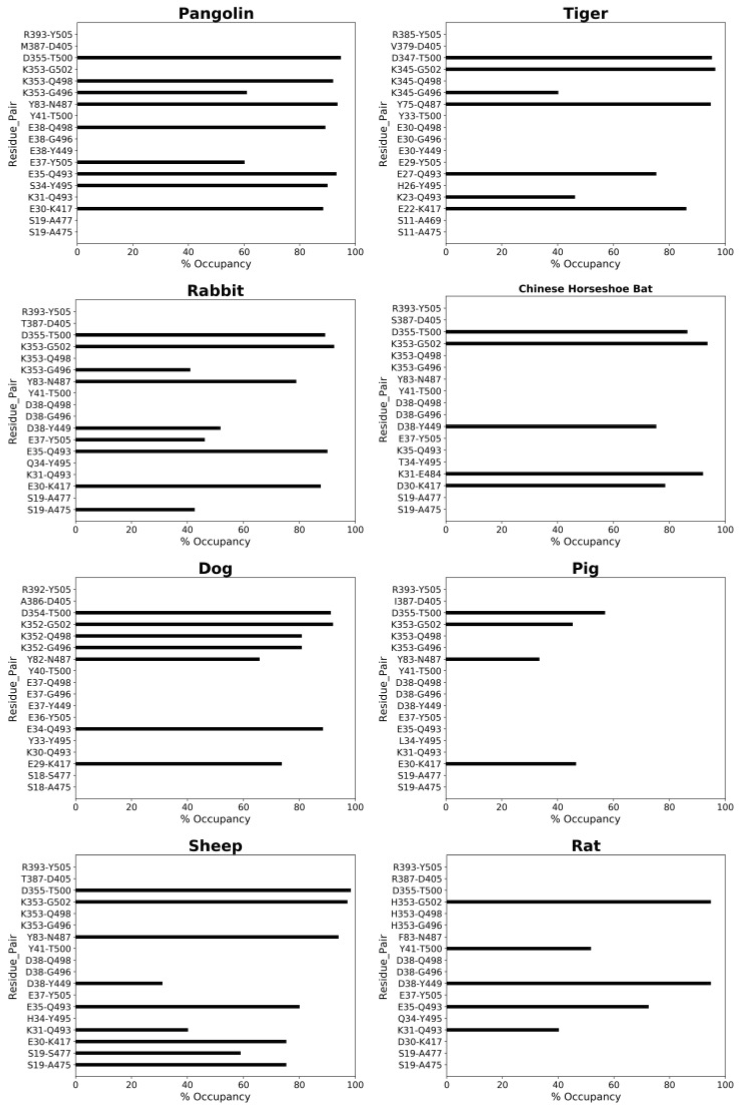

**Figure S5**. Occupancies of H-bonds at ACE2-RBD interface for eight different ACE2 proteins. Each residue pair comprises a residue from ACE2 (left of the hyphen) and a residue from spike RBD (right of the hyphen).
